# Monitoring *S. pombe* genome stress by visualizing end-binding protein Ku

**DOI:** 10.1101/2020.06.12.149054

**Authors:** Chance Jones, Susan L Forsburg

## Abstract

Studies of genome stability have exploited visualization of fluorescently tagged proteins in live cells to characterize DNA damage, checkpoint, and repair responses. In this report, we describe a new tool for fission yeast, a tagged version of the end-binding protein Pku70 which is part of the KU protein complex. We compare Pku70 localization to other markers upon treatment to various genotoxins, and identify a unique pattern of distribution. Pku70 provides a new tool to define and characterize DNA lesions and the repair response.

## Introduction

The response to genome stress and DNA repair can be observed in living cells in real time, by monitoring fluorescently-tagged DNA damage response proteins (Lisby, M. *et* al. 2004; Lukas, C., *et* al. 2005; Nagy, Z., & Soutoglou, E., 2009; Polo, S. E., & Jackson, S. P. 2011). This has allowed characterization of dynamic processes that respond to damage and preserve genome integrity, including cell cycle, checkpoint, repair, and recovery pathways. In the fission yeast *S. pombe*, accumulation of foci of the single-strand DNA binding protein Ssb1 (a subunit of Replication Protein A /RPA), and of the recombination protein Rad52, have been used to characterize intrinsic genome stresses as well as the response to external genotoxins (Meister, P., *et* al. 2003; Kilkenny, M. L., *et* al. 2008; Carneiro, T., *et* al. 2010; Bass, K. L., *et* al 2012; Sabatinos, S.A., *et* al. 2012). These proteins recognize and respond to single strand DNA accumulation, which can result from exonuclease activity, resection, processing of replication forks and recombination intermediates, or R-loop or D-loop formation (Zeman, M. K., & Cimprich, K. A., 2014)., Sabatinos, S. A., & Forsburg, S. L., 2015). Importantly, this has led to identification of distinct patterns of accumulation that can serve as fingerprints for different forms of genome stress (e.g., Sabatinos, S.A., *et* al. 2012; Sabatinos, S.A., *et* al. 2015 Ranatunga).

The fission Pku70 protein is the orthologue of the Ku70 subunit of the conserved heterodimeric Ku complex (Baumann, P., & Cech, T. R., 2000). Ku is abundant and binds efficiently to DNA double strand breaks (DSBs) (Fell, V. L., & Schild-Poulter, C. 2015; Shibata, A., *et* al. 2018). Ku is associated with the non-homologous end-joining (NHEJ) mechanism of DNA double strand break (DSB) repair (Mahaney, B. L., *et* al. 2009) and protects telomeres (Baumann, P., & Cech, T. R., 2000; Ferreira, M. G., & Cooper, J. P. 2001). Additionally, it recognizes “one-sided” double strand breaks and ends associated with regressed replication forks (Teixeira-Silva, A., *et* al. 2017; Foster, S. S., *et* al. 2011; Langerak, P., *et* al. 2011).

Ku binding to DNA ends inhibits the resection and accumulation of single strand DNA that otherwise drives homologous recombination (Shibata, A., *et* al. 2018). Its activity is coordinated with the Mre11-Rad50-Nbs1 (MRN) protein complex, another early responder to DNA double strand breaks (Shibata, A., *et* al. 2018; Syed, A., & Tainer, J. A. 2018). MRN is also linked to DNA DSB end binding (Wang, Q., *et* al. 2014) and resection (Shibata, A., *et* al. 2014) and contributes to DNA damage checkpoint activation (Chahwan, C., *et* al. 2003; Paull, T. T. 2015). The Mre11/Rad32 subunit is able to drive endonucleolytic cleavage of DNA ends that are blocked by covalently bound proteins such as Spo11 or Top2 (Hartsuiker, E., *et* al. 2009; Milman, N., *et* al. 2009; Rothenberg, M., *et* al. 2009; Hartsuiker, E., *et* al. 2009; Garcia, V., *et* al. 2011; Reginato, G., *et* al. 2017). To some degree, Ku and MRN act as mutual antagonists; Ku inhibits short-range resection driven by MRN, and MRN removes Ku to facilitate homologous recombination (HR) over NHEJ; and to prevent inappropriate repair of single-end breaks (Langerak, P., *et* al. 2011; Shao, Z., *et* al. 2012; Myler, L. R., *et* al. 2017; Shibata, A., *et* al. 2018). Interestingly, loss of Ku partly suppresses the sensitivity to DNA damage and replication blocking toxins associated with mutation of MRN (Tomita, K., *et* al. 2003; Williams, R. S., *et* al. 2008; Limbo, O., *et* al. 2007; Langerak, P., *et* al. 2011; Teixeira-Silva, A., *et* al. 2017), which can lead to excessive Exo1 driven resection but impaired RPA recruitment (Teixeira-Silva, A., *et* al. 2017).

In this report, we describe the development of a new fluorescent marker for fission yeast, the Pku70 subunit of the Ku protein complex that recognizes DNA ends. We constructed a *pku70*^*+*^ -citrine fusion and integrated into the genome in wild type fission yeast under the endogenous promoter. We examined its behavior and accumulation in treated and untreated wild type cells in response to different genotoxins. We compared localization of Ku to Rad52, RPA, and Mre11 markers and observe a pattern of foci that is distinct from other markers. This provides a new tool to characterize responses to different forms of genotoxic stress and a useful addition to the fission yeast tool kit for investigation of the 3-Rs of DNA replication, repair, and recombination.

## Results

### Construction of strains with fluorescently tagged Pku70 and Mre11

Ku (a heterodimer of Pku70/80) and MRN (Mre11/Rad50/Nbs1) protein complexes are known for high affinity for binding DNA ends (Fell, V. L., & Schild-Poulter, C. 2015; Shibata, A., *et* al. 2018). We tagged Pku70 on its C-terminal end with Citrine fluorescent protein and integrated into the endogenous locus (see methods). Using a similar strategy we also tagged Mre11 on its C terminal end with mCherry fluorescent protein. The resulting strains were compared to wild type, *pku70Δ, mre11Δ* and *rad51Δ* for their growth on four typical genotoxic drugs: methyl methanesulfonate (MMS), which creates alkylation damage that inhibits DNA replication fork progression; camptothecin (CPT), which blocks Topoisomerase I cleavage; hydroxyurea (HU), which causes nucleotide starvation and fork pausing; and Phleomycin (phleo), a radio-mimetic that causes single- and double-strand breaks. Both the Mre11-mCherry and Pku70-Citrine tagged strains behaved the same as WT under normal growth and genotoxic stress. The *Δpku70* strain also shows no sign of genotoxin sensitivity, as reported previously (Manolis, K. G., *et* al. 2001; Sánchez, A., & Russell, P. 2015). (Fig. 1A)

**Figure 1.**
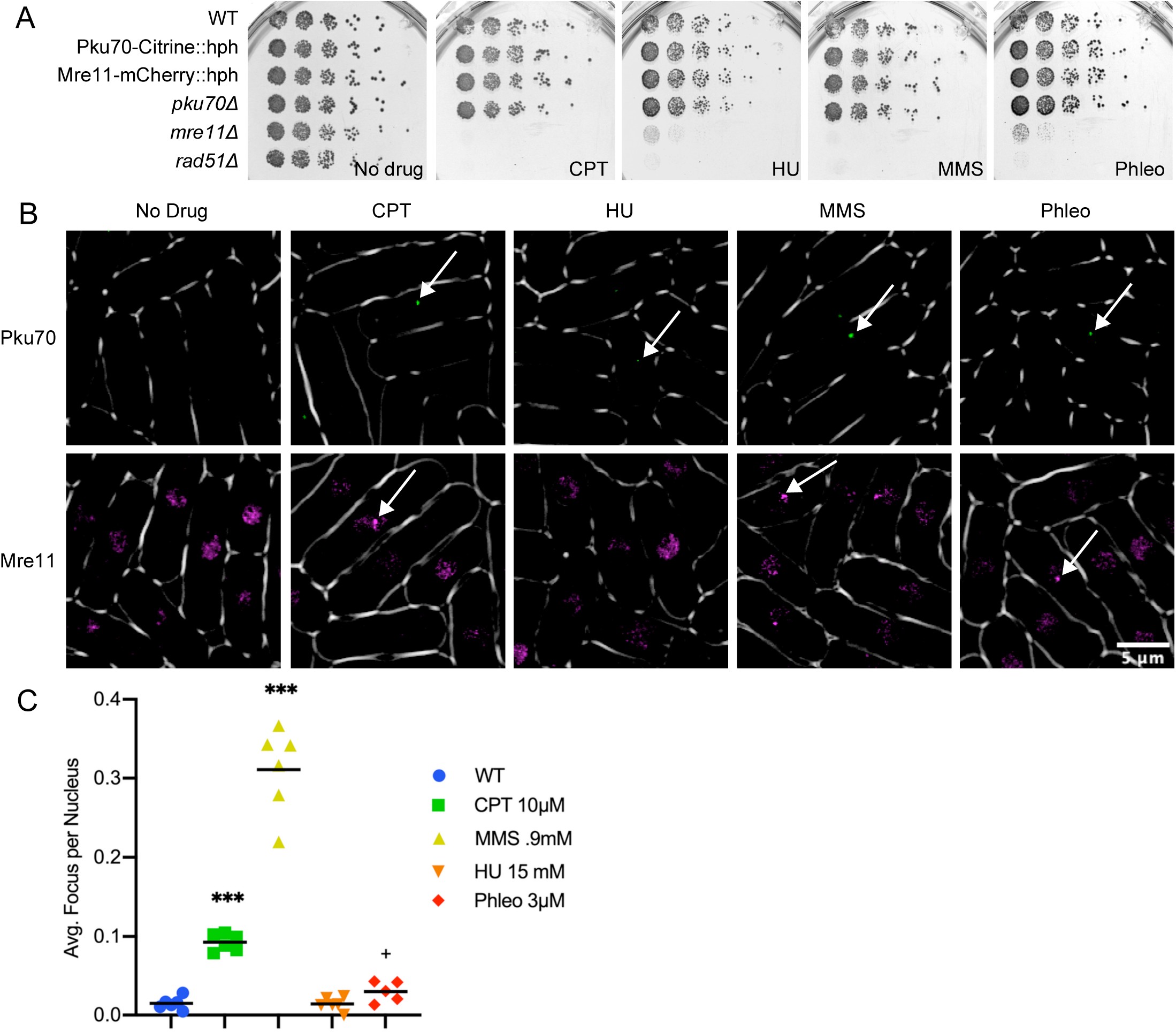
Construction of Fluorescently Tagged Strains. A) Frogging of flurescently tagged strains on various genotoxic drugs. Drug concentrations are as follows, CPT .0125 mM, HU 7.5 mM, MMS .012%, Phleo 1.5 mU. B) Fluorescently tagged strains Pku70-Citrine::hph and Mre11-mCherry::hph both with and without genotoxic drugs. For clarity Mre11-mCherry is shown in false color as magenta. Arrows show representative foci for Pku70 and generalized areas of increased fluorescence for Mre11. B C) Pku-Citrine foci were counted by hand and significance testing was done using a Mann Whitney significance test. Six replicates were used per drug tested.

### Pku70 and Mre11 have increased nuclear signal following genotoxic stress

When we imaged the tagged strains under normal growth conditions, we observed a few scattered foci in Pku70-citrine cells and diffuse nuclear fluorescence in Mre11-mCherry cells (Fig. 1B). We next examined the distribution of signal in cells treated with MMS, CPT, Phleo, or HU at 32°C after 4 hours. There is a significant increase of cells with individual Pku70 nuclear foci in MMS, CPT, and to a lesser extent Phleo. Cells treated with HU did not show any significant difference from WT (Fig. 1C). In contrast, the Mre11-mCherry signal showed diffuse pan-nuclear staining in untreated cells (Fig 1B). Following 4 hours of treatment with the four genotoxic drugs, Mre11-mCherry did not show obvious foci. Rather, we observed generalized areas of increased fluorescence over threshold, but these typically were not well-defined discrete puncta as seen with other markers.

### Colocalization of Pku70 and Mre11 with other markers of DNA damage

Previous studies of genome instability in fission yeast have imaged the single stranded binding protein Ssb1 (Rad11, RPA) and the homologous recombination protein Rad52 in response to different forms of replication stress (Meister, P., *et* al. 2003; Kilkenny, M. L., *et* al 2008; Carneiro, T., *et* al. 2010; Bass, K. L., *et* al., 2012; Sabatinos, S. A., *et* al. 2012). We examined co-localization using CPT, MMS, HU and Phleo in a strain with Pku70-citrine, Rad52-mCherry, and RPA-CFP. Four hours after drug addition at 32°C, we determined frequency of colocalization among all three tagged proteins. While there was partial overlap, Ku is not completely concordant with the other markers (Fig. 2A).

**Figure 2.**
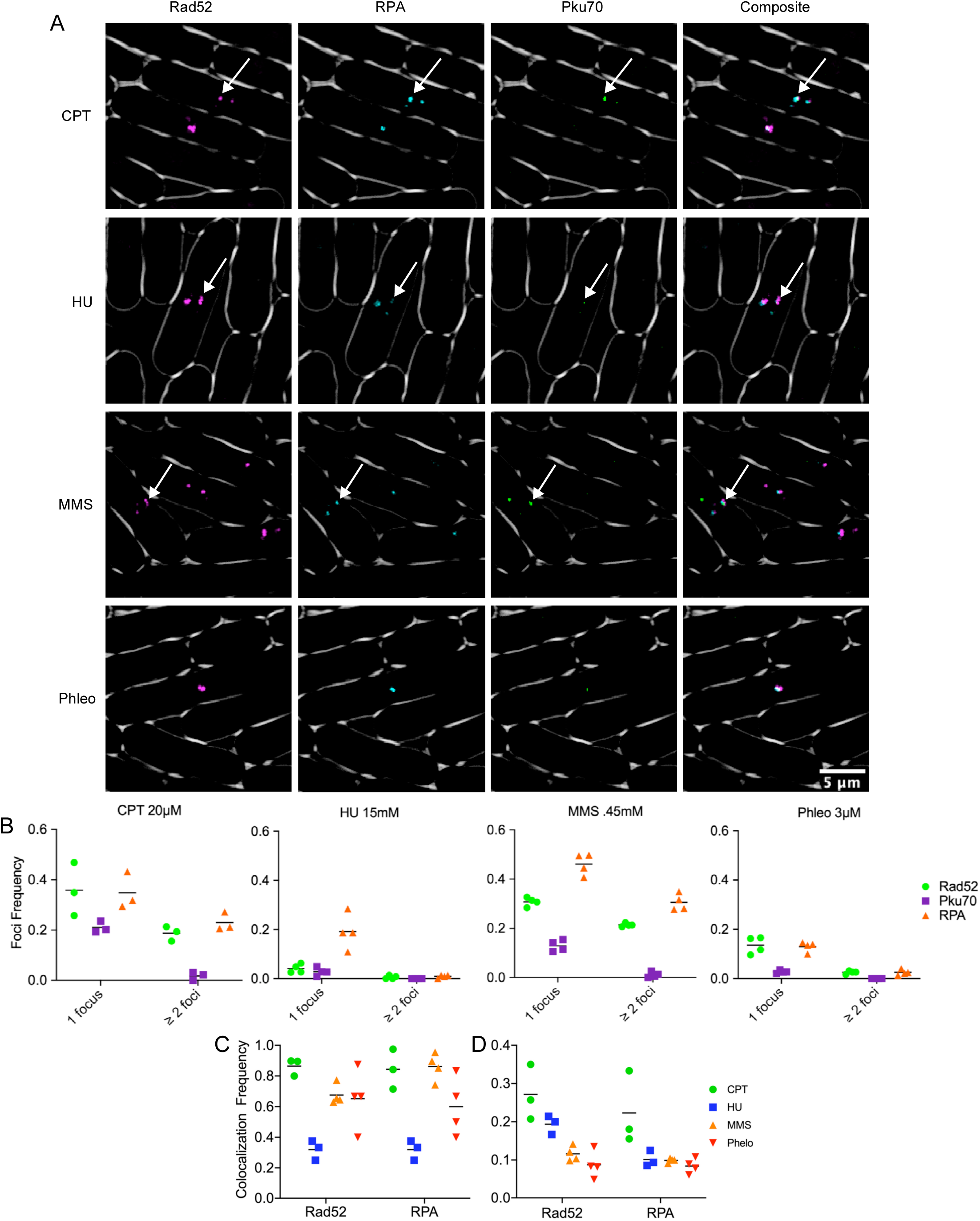
Pku70-Citrine colocalization with DNA repair proteins. A) These images depict colocalization of Pku70-Citrine with previously reported HR repair proteins Rad52-mCherry and RPA-CFP under four commonly used genotoxic drugs. For clarity Rad52-mCherry is shown in false color magenta, Pku70-Citrine is being shown in false color as green. In the composite image overlapping foci appear as white. Arrows show examples of easily visible colocalization. B) Percent of nuclei with either 1 or ≥2 Pku70-citrine foci. C) Percent of Pku70 foci that colocalize with either a Rad52 foci or a RPA foci. D) Percent of either Rad52 or RPA foci that have a corresponding colocalizing Pku70 foci.

The number of foci per nucleus was calculated and binned as either 1or ≥ 2 foci using an automatic foci counter in ImageJ as described in the materials and methods (Fig. 2B). We observed that CPT 20µM contained the highest frequency of Pku70 foci, then MMS, Phleo, and HU. The difference from prior observation likely reflects a somewhat different drug dosage: CPT levels were raised from 10µM to 20 µM in order to produce an enhanced response and MMS was lowered from .9mM to .45mM to better resolve single foci.

Colocalization was determined using the objects-based method in the ImageJ pluggin JACoP (see materials and methods). Fig. 2C shows the proportion of Pku70-Citrine foci that overlap with a thresholded region for Rad52-mCherry or RPA-CFP. For CPT, MMS, and Phleo these proportions vary from 60-90%. In contrast, the scattered foci in HU showed only about 30% of Ku co-associating with another marker. Fig 2D shows the proportion of Rad52-mCherry foci that have a colocalizing Pku70-Citrine focus. CPT contained the highest proportion of Rad52 as well as RPA with overlapping Pku70 foci, whereas HU contained the lowest.

We performed a similar study with Mre11-mCherry but could not perform the same quantitation because Mre11-mCherry does not form discrete foci. We observed areas of generally increased fluorescence but never clear puncta as with Pku70, Rad52, or RPA. Observing these cells in three-dimensional reconstruction showed no obvious colocalization between Rad52-YFP and RPA-CFP in live cell video microscopy, or in static images (Supplemental Fig. 1; Fig. 3A,B).

**Figure 3.**
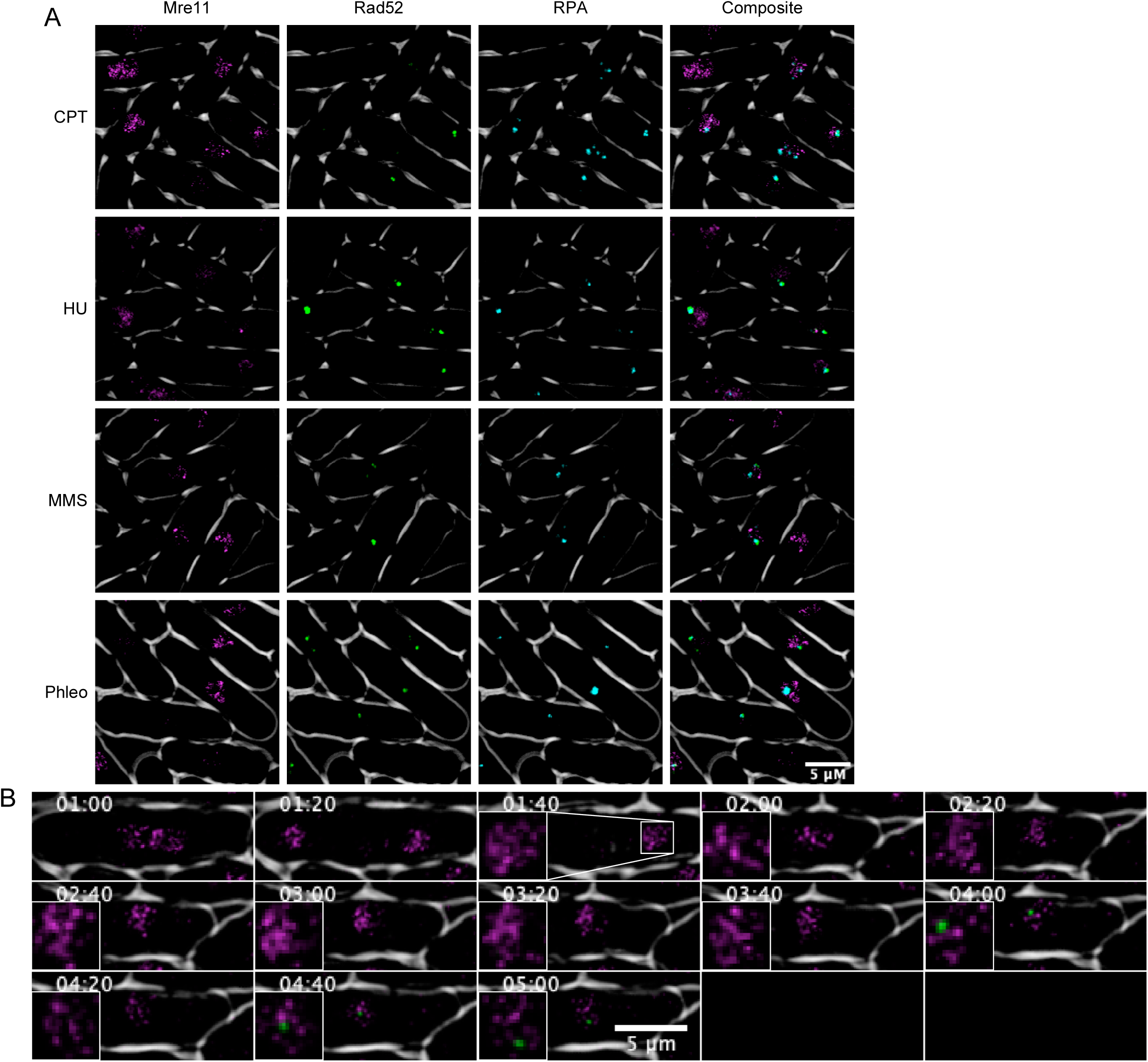
Colocalization of Mre11-mCherry, Rad52-YFP, RPA-CFP and Pku70-Citrine. **A)** Cells were treated in .45mM MMS for 4 hours at 32°C. Mre11-mCherry is shown in false color as magenta and Rad52-YFP is shown in green for clarity. B) Timelapse microscopy of Mre11-mCherry and Pku70-Citrine. Cells were treated in .45mM MMS and time-lapses were kept 28°C. Timepoints designate time since drug addition.

### Pku dynamics in S phase specific damage

The genotoxin MMS causes alkylation damage, generating lesions that block DNA polymerase (Lundin, C., *et* al. 2005). This typically results in replication template switching (Barbour and Xiao 2003; Andersen et al. 2008). Previous work has suggested that Ku is recruited by blocked and regressed replication forks (Teixeira-Silva, A., *et* al. 2017). Therefore, we investigated the dynamics of Ku response to MMS treatment as a model for disruptions in fork progression. We used live cell video microscopy to observe cells containing Rad52-mCherry and Pku70-Citrine over a 5hr period of MMS treatment at 28°C. We observe distinct dynamics for Rad52 and Pku70 recruitment during treatment. While absolute timing differs in individual cells, typically a Ku focus appears for a short time and partially co-localized with Rad52.

Fig 4A shows a representative newborn cell that has just entered S phase, 1h and 20m after drug treatment. The diffuse Rad52-mCherry signal distributes into smaller foci which then coalesce into two large foci. Pku70-Citrine colocalizes at the center of these large foci for about 20-40 minutes. The large Rad52-mCherry foci persist another 60 minutes and then begin to dissipate. Retention time of Pku-Citrine foci in MMS is ≤ 20 minutes with a fraction of cells maintaining it longer between 20 and 40 minutes. In contrast, Rad52 foci extend over a much longer period of time ranging from 20 all the way up to 160 minutes (Fig. 4B). Overall Rad52 tends to appear slightly earlier than Pku70 in most cells and disappears much later (Fig. 4C). (Additional time-lapse images found in Supplemental Fig. 2 and 3)

**Figure 4.**
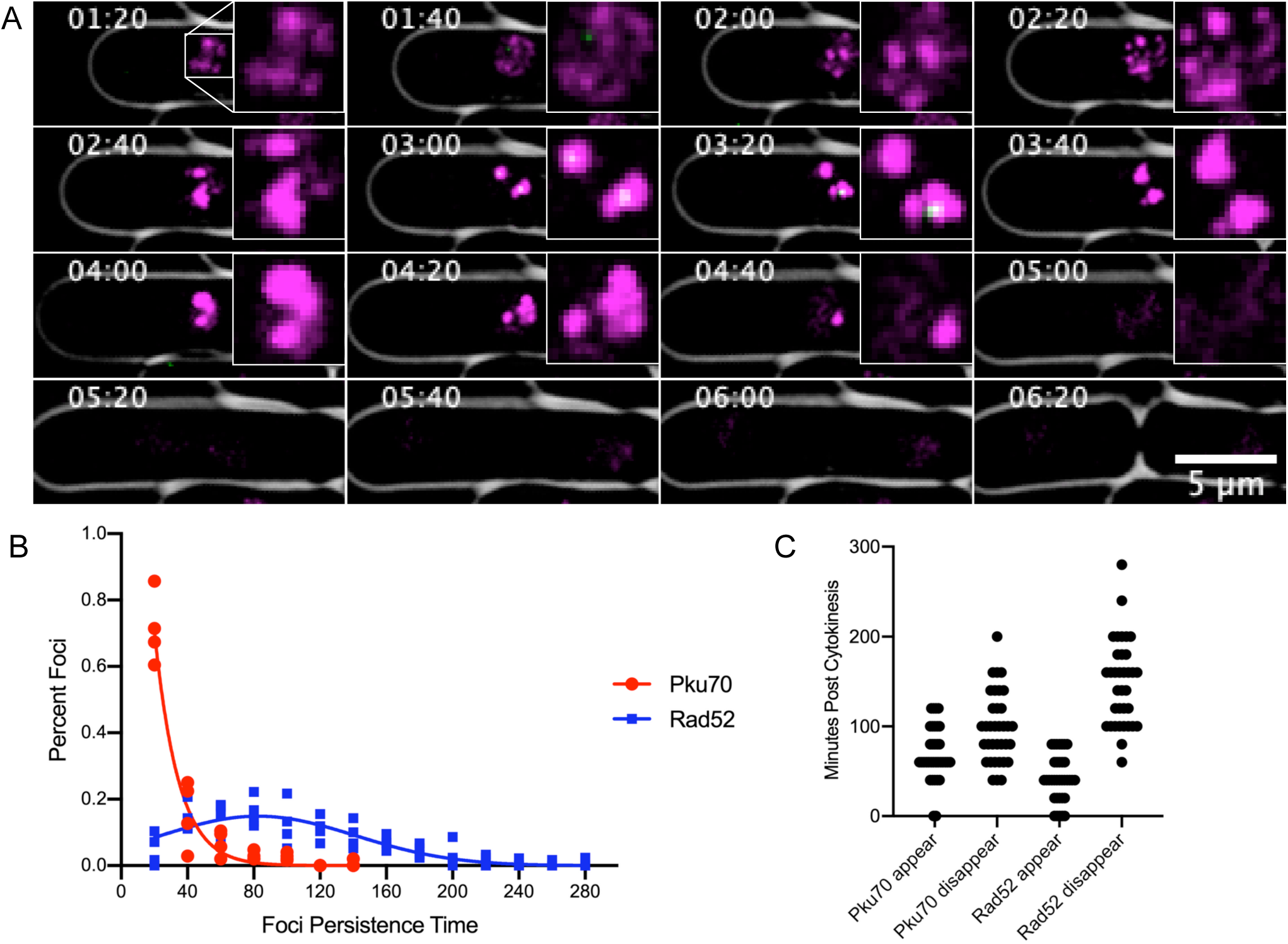
Pku localization in a dynamic timecourse of MMS treatment. A) Fluorescent time lapse images of Pku70-Citrine colocalizing with Rad52-mcherry. For clarity Rad52-mCherry is shown in magenta and Pku-Citrine is shown in green. Imaging was started at 80 minutes post addition of .45mM MMS and cells were imaged at 28°C. Timecourse images were taken every 20 min. B) Persistence time**ds** of Rad52-mCherry (n=205) and Pku70-Citrine (n=195). C) Appearance and dissapearance times for Pku70-Citrine and Rad52-YFP in in**F** dividual mononucleate cells. T=0 first time point after completion of cytokinesis. Horizontal density shows higher quantity of foci appearing or dissapearing at that time point. (n=35 cells).

## Discussion

Localization of repair puncta in fission yeast has been a well-established means of observing DNA damage, quantified by counting foci, determining pixel intensity or size of foci, and three-dimensional position in the nucleus (Green, M. D., *et* al., 2015). The most frequently used fluorescent tags used in *S. pombe* for observing DNA lesions are the recombination protein Rad52 and single strand DNA binding protein Rad11, a subunit of RPA (Meister, P., *et* al., 2003; Carneiro, T., *et* al. 2010; Sabatinos, S. A., *et* al. 2012). Studies have shown that in cycling wild type cells, approximately 10-20% of cells show evidence of single RPA or Rad52 foci, likely due to sporadic S phase events. These proteins show distinct patterns in response to genotoxic stresses induced by mutations in the replication or repair pathways (Sabatinos, S. A., *et* al. 2012; Sabatinos, S. A., *et* al. 2015; Ranatunga, N. S., & Forsburg, S. L. 2016), or in response to exogenous agents such as hydroxyurea (HU), which causes replication fork stalling (Thelander, L., & Reichard, P. 1979); MMS, an alkylating agent that generates lesions that block the replication fork (Lundin, C., *et* al. 2005); camptothecin (CPT), a topoisomerase I inhibitor that leads to S-phase specific double strand breaks (Li, T. K., & Liu, L. F. 2001); and bleo- or phleomycin, radiomimetic drugs that causes single- and double-strand breaks (Povirk, L. F. 1996).

The current study seeks to expand the library of tagged proteins, part of our strategy to develop a fingerprint for the response to different forms of genotoxic stress. We investigated fluorescently tagged Mre11 and Pku70 as markers for DNA breaks.

The MRN complex is one of the earliest responders to DSBs (Shibata, A., *et* al. 2018; Syed, A., & Tainer, J. A. 2018). and is essential to drive resection (Wang, Q., *et* al. 2014; Shibata, A., *et* al. 2018; Langerak, P., *et* al. 2011; Teixeira-Silva, A., *et* al. 2017). Our Mre11-mCherry construct showed a diffuse pan-nuclear signal in untreated cells. We did not see obvious focus formation of Mre11-mCherry following treatment with genotoxins. Rather, it maintained a diffuse signal with regions of brightness. In other systems, MRN has been shown to be an immediate responder to double strand breaks induced by ionizing radiation (Maser, R. S., *et* al. 1997). Our failure to see this form of localization may indicate the timing of our analysis, and diffuse distribution of lesions in drug treated cells, compared to concentrated sites of damage from of ionizing radiation.

Previous whole-cell localization of Pku70 in S. pombe was carried out using C terminal epitope tagged Pku70 and immunofluorescence on fixed cells (Manolis, K. G., *et al*. 2001). In unperturbed cells, there is a diffuse pan-nuclear localization. Association of Ku with DNA ends has been investigated using ChIP methods; in wild type cells, it is not enriched unless the MRN complex is missing (Langerak, P., *et* al. 2011; Teixeira-Silva, A., *et* al. 2017). Visualization of Pku70 in live fission yeast cells has not previously been performed.

We saw few Ku foci in WT cells, consistent with previous reports. Treatment for 4 hours with our panel of genotoxins showed that HU has little to no accumulation of Ku foci. Treatment with CPT causes a modest but limited increase in the fraction of cells with foci at 10µM and a much more drastic increase at 20µM. Similarly, phleomycin, a radiomimetic that causes DNA breaks throughout the cell cycle, has a modest but limited increase in foci.

The most dramatic increase in fraction of cells with foci was observed with MMS at 0.9mM, an alkylating agent that results in error-free and error prone base excision repair during S phase, and thus leading to trans lesion synthesis. This induction in MMS is consistent with prior observations suggesting that Ku is recruited to regressed or broken replication forks in order to stabilize the free end (Langerak, P., *et* al. 2011; Teixeira-Silva, A., *et* al. 2017). This suggests that even in MRN^+^ cells, there are situations where Ku remains associated with sites of genome stress.

We observed a substantial colocalization between RPA or Rad52 and Ku, in cells treated with MMS, CPT, or Phleo. This result was a surprise as many models suggest Pku should be removed by the time resection and recombination proteins are recruited. One possibility for the S phase specific toxins is that Pku could be binding to reversed forks at repair centers. Previous studies suggest that Pku plays a role at reversed forks in order to maintain genome stability, particularly in cells with defective HR repair such as *brc1Δ* (Sánchez, A., & Russell, P. 2015; Teixeira-Silva, A., *et* al. 2017). This may reflect that other mechanisms generate ssDNA besides exonuclease activity, including helicase unwinding and strand invasion.

To address this finding in dynamic conditions, we examined MMS-treated cells as a model for stalled replication forks. Previously, we showed that MMS induces a dramatic increase in RPA and Rad52 foci relative to other genotoxins (Ranatunga, N. S., & Forsburg, S. L. 2016). We observe substantial recruitment of Rad52-mCherry and brief, partial co-localization of Pku70. The Pku70 signal, largely in 1-2 foci, appears after Rad52 and disappears before Rad52 is resolved. Further molecular work will be required to determine what this signal represents.

It is likely that Ku foci will define distinct structures associated with particular forms of replication stress. For example, in a recent study, we showed that a mutant *mcm4-dg* with a defect in the MCM helicase accumulates Ku foci (Kim, S. M., & Forsburg, S. L. 2020). This accumulation can be reversed by activation of the Mus81 resolvase. Mus81 is essential for viability in *pku80Δ brc1Δ* mutants (Sánchez, A., & Russell, P. 2015), indicating a collaboration between Ku and Mus81 in response to replication stress. Our Pku70-citrine fusion will be a key reagent in dissecting this and other activities.

## Methods

### Cell growth and physiology

Fission yeast strains are described in Table 1, and were grown as in (Sabatinos *et* al., 2012).

**Table 1:** Strains

|  |  |  |
| --- | --- | --- |
| FY527 | h- his3-D1 ade6-M216 ura4-D18 leu1-32 | Gould, K. L., 1998 |
| FY528 | h+ his3-D1 ade6-M210 ura4-D18 leu1-32 | Liang, D. T., 1999 |
| FY8488 | h+ Pku70-citrine::hph his3-D1 ade6-M210 ura4-D18 leu1-32 | This study |
| FY8558 | h- Pku70-citrine::hph his3-D1 ade6? ura4-D18 leu1-32 | This study |
| FY8661 | h+ Mre11-mCherry::hph his3-D1 ade6-M210 ura4-D18 leu1-32 | This study |
| FY8662 | h- Mre11-mCherry::hph his3-D1 ade6? ura4-D18 leu1-32 | This study |
| FY8381 | h- Rad52-mCherry::kan ura4-D18 leu1-32 | Yu, Y., 2013 |
| FY8625 | h- Pku70-citrine::hph Rad52-mCherry::kan his3-D1? ade6? ura4-D18 leu1-32 | This study |
| FY8698 | h90 Mre11-mCherry::hph, Pku70-citrine::hph, his3-D1 ade6? ura4-D18 leu1-32 | This study |
| FY4743 | h- rad11-Cerulean::hphMX rad22-YFP-natMX leu1-32 ade6-M210 ura4-D18 | Sabatinos, S. A., 2012 |
| FY8687 | h90 Mre11-mCherry::hph, RPA-Cerulean::hphMX, Rad52-YFP-natMX his-D1? ade6-M210 ura4-D18 leu1-32 | This study |
| FY9381 | h- Rad11-Cerulean::hphMX pKu70-citrine::hph rad52-mCherry::natMX6 ura4-D18 leu1-32 his? ade? | This study |

**Table 2:**
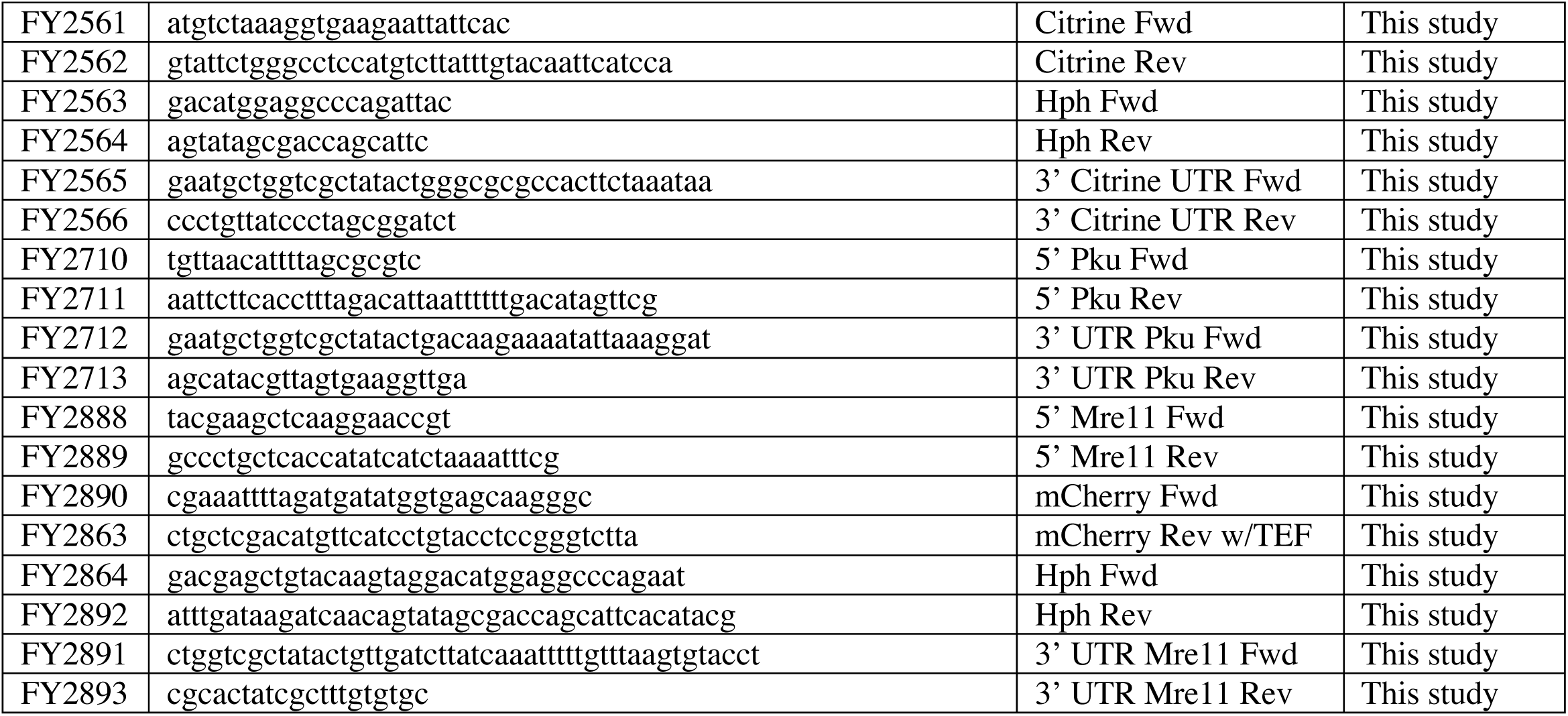
Primer List

### Construction of Tagged strains

All fragments were lengthened using the Expand Long Template PCR System (Roche Diagnostics, Mannheim Germany). Primers were designed using the NCBI Primer design tool and optimized to an annealing temperature of 52-54°C (Ye J, *et* al. 2012). Full length fragments were transformed using electroporation and selected using the appropriate marker (Sabatinos, S. A., & Forsburg, S. L. 2010). Upon transformation instead of plating directly onto selective minimal media the cells were first plated on YES for 24 hours then replica plated onto YES-Hph. Candidate colonies growing on Hph after 4-5 days were then restreaked onto Hph twice and visually screened for nuclear localizing foci.

### Pku-Citrine::Hph

The Pku C-terminal Citrine fragment was formed from 5 fragments, Pku 5’overhand (FY2710 + FY2711), Citrine (FY2561 + FY2562), Hph (FY2563 + FY2564), Citrine UTR (FY2565 + FY2566), and Pku 3’ UTR overhang (FY2712 + FY2713). The Citrine and Citrine UTR fragments were lengthened from addgene plasmid pKT0139 (Sheff, M. A., & Thorn, K. S. 2004). The Hph fragment was lengthened from pFA6a-hphMX6 (Hentges, P., *et* al. 2005). The 5’ and 3’ UTR overhang fragments were lengthened from phenol:chloroform extracted WT (FY527) DNA (Forsburg, S. L., & Rhind, N. 2006). The Citrine, Hph, and Citrine UTR fragments were first lengthened to form a full Citrine::HPH fragment. A single PCR reaction was then done with Pku 5’overhang, Citrine HPH, and Pku 3’ UTR overhang fragments forming the full fragment. This fragment was then used for transformation.

### Mre11-mCherry::Hph

The Mre11 C-terminal mCherry fragment was formed from 4 fragments, Mre11 5’ overhang (FY2888 + 2998), mCherry (FY2890 + FY2863), Hph (FY2864 + FY2892), Mre11 3’ UTR overhang (FY2891 + FY2893). The mCherry fragment was lengthened from extracted DNA, FY8381 (Yu, Y., *et* al. 2013). The Hph fragment was lengthened from the previously formed Citrine::Hph fragment above. The Mre11 5’ and 3’ UTR overhang fragments were lengthened from extracted WT DNA (FY527). The mCherry::Hph fragment was first lengthened. The Mre11 5’ overhang, mCherry::Hph, and Mre11 3’ UTR overhang fragments were then combined in one PCR reaction forming the full fragment. The fragment was then used for transformation.

### Live Cell Imaging

Cells were prepared as in (Green, M. D., *et* al. 2015) Medium for all live cell imaging was PMG-HULALA (PMG + Histidine, Uracil, Leucine, Adenine, Lysine, Arginine) (225mg/L each) (Sabatinos, S. A., & Forsburg, S. L. 2010) Unless specified all drug concentrations used for imaging were as follows, MMS .9mM, HU 15mM, CPT 20µM, Phleo 3µM. Strains in liquid cultures at 32°C were grown to mid-log phase. Cells concentrated by a brief microfuge spin were applied to 2% agarose pads made from PMG + HULA and prepared on glass slides sealed with VaLaP (1/1/1 w/w/w vasoline/lanolin/paraffin). Static images were performed at room temperature 22°C and long term timelapse images were taken at a constant temperature of 28°C. Images were acquired with a DeltaVision Core (Applied Precision, Issaquah, WA) microscope using a 60x N.A. 1.4 PlanApo objective lens and a 12-bit Photometrics CoolSnap HQII CCD. The system x-y pixel size is 0.109µm. softWoRx v4.1 (Applied Precision, Issaquah, WA) software was used at acquisition. Excitation illumination was from a Solid-state illuminator, CFP was excited and detected with a 438/24,470/24 filter set (excitation intensity attenuated to 10%) and a 400ms exposure; YFP was excited and detected with a 513/17,559/38 (excitation intensity attenuated to 32% for Rad52-YFP and 50% for Pku70-Citrine) filter set and a 200ms exposure. A suitable polychroic mirror was used. Sections of static timepoints were 20 .20µm z-sections. Long-term time-lapse videos used 8 z-steps of .35µm. 3-D stacks were deconvolved with manufacturer provided OTFs using a constrained iterative algorithm, images were maximum intensity projected for presentation. Images were contrast adjusted using a histogram stretch with an equivalent scale and gamma for comparability. Brightfield images were also acquired.

### Image processing and analysis

Images were contrast adjusted using an equivalent histogram stretch on all samples. Significance was assessed with Mann Whitney tests. For publication Long-term time lapse videos were stabilized in ImageJ using the package “StackReg” by Philippe Thevanaz from the Biomedical Imaging Group at the Swiss Federal Insitute of Technology Lausanne. (Thevenaz, P., *et*. Al 1998). Foci were automatically quantified using a computational algorithm based on uniform threshold per fluorescence channel as described by the light microscopy core facility at Duke University (https://microscopy.duke.edu/guides/count-nuclear-foci-ImageJ. Object based colocalization analysis was performed using the ImageJ pluggin JACoP on the same images used for the focu quantification. However this object based colocalization analysis method still requires observer-based thresholding before analysis. In order to mitigate observer-based thresholding bias the number of observed objects after thresholding per fluorescence channel was calculated to be within 10 foci of the automatically counted foci during the previous computer-based foci quantification analysis described above.

---

**Supplemental Figure 3.**
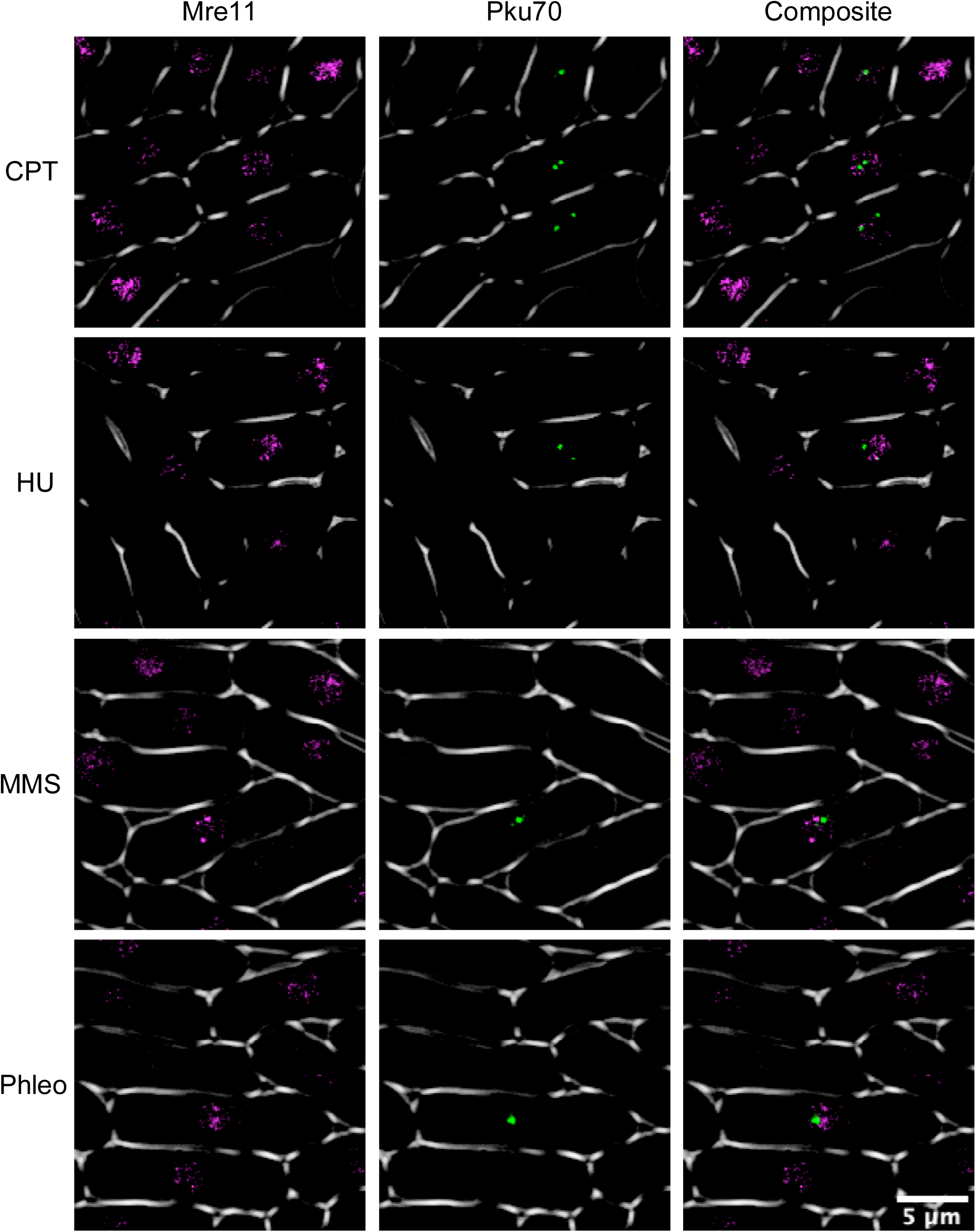
Colocalization of Pku70 and Mre11. Cells were treated in .45mM MMS for 4 hours at 32°C. Mre11-mCherry is shown in false color as magenta and Pku70 is shown in green for clarity.

**Supplemental Figure 2.**
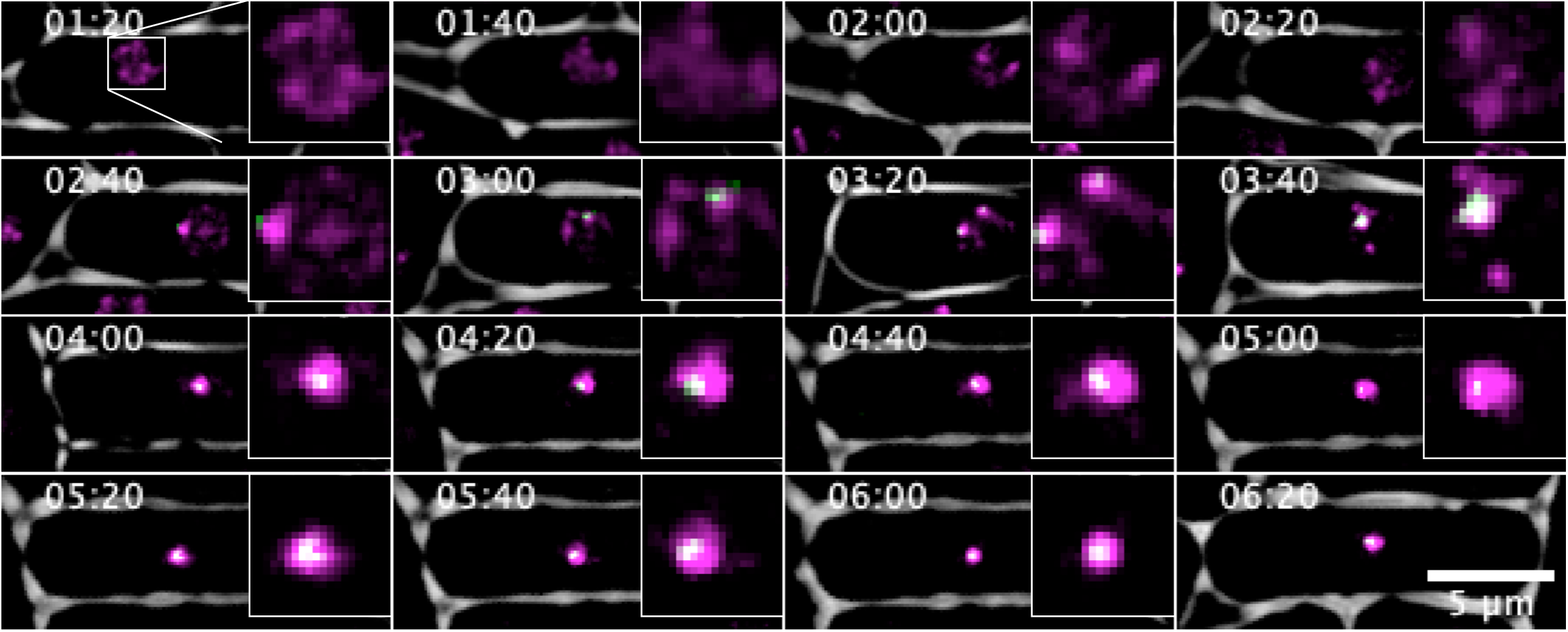
Timelapse of Pku70 and Rad52. Fluorescent time lapse images of Pku70-Citrine colocalizing with Rad52-mcherry. For clarity Rad52-mCherry is shown in magenta as well as Pku-Citrine being shown in green. Imaging was started at 80 minutes post addition of .45mM MMS and cells were grown at 28°C. Timecourse images were taken every 20 min.

**Supplemental Figure 3.**
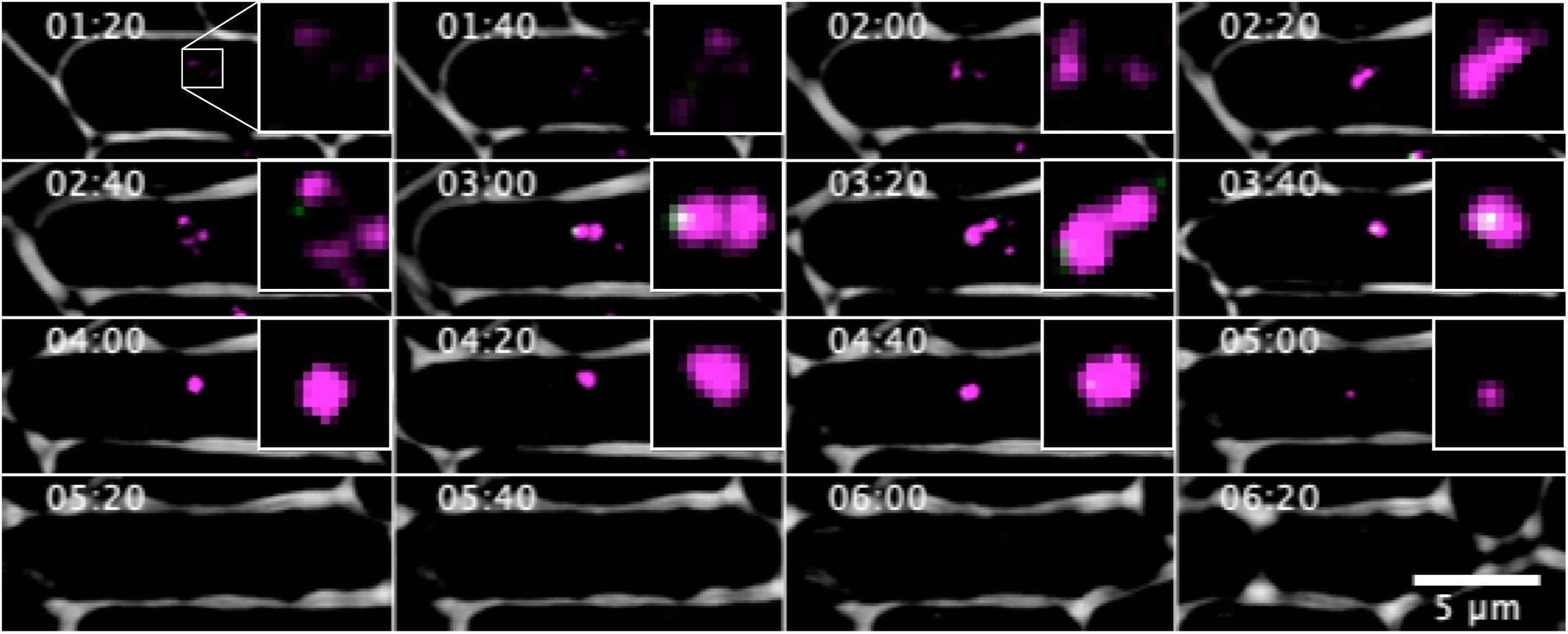
Timelapse of Pku70 and Rad52. Fluorescent time lapse images of Pku70-Citrine colocalizing with Rad52-mcherry. For clarity Rad52-mCherry is shown in magenta as well as Pku-Citrine being shown in green. Imaging was started at 80 minutes post addition of .45mM MMS and cells were grown at 28°C. Timecourse images were taken every 20 min.

